# Sign-tracking behavior is difficult to extinguish and resistant to multiple cognitive enhancers

**DOI:** 10.1101/581769

**Authors:** Christopher J. Fitzpatrick, Trevor Geary, Justin F. Creeden, Jonathan D. Morrow

**Author notes:** Contact Information: Dr. Jonathan Morrow, 5047 Biomedical Sciences Research Building, 109 Zina Pitcher Place, Ann Arbor, MI, USA 48109.

## Abstract

The attribution of incentive-motivational value to drug-related cues underlies relapse and craving in drug addiction. One method of addiction treatment, cue-exposure therapy, utilizes repeated presentations of drug-related cues in the absence of drug (i.e., extinction learning); however, its efficacy has been limited due to an incomplete understanding of extinction and relapse processes after cues have been imbued with incentive-motivational value. To investigate this, we used a Pavlovian conditioned approach procedure to screen for rats that attribute incentive-motivational value to reward-related cues (sign-trackers; STs) or those that do not (goal-trackers; GTs). In Experiment 1, rats underwent Pavlovian extinction followed by reinstatement and spontaneous recovery tests. For comparison, a separate group of rats underwent PCA training followed by operant conditioning, extinction, and tests of reinstatement and spontaneous recovery. In Experiment 2, three cognitive enhancers (sodium butyrate, D-cycloserine, and fibroblast growth factor 2) were administered following extinction training to facilitate extinction learning. STs but not GTs displayed enduring resistance to Pavlovian, but not operant, extinction and were more susceptible to spontaneous recovery. In addition, none of the cognitive enhancers tested affected extinction learning. These results expand our understanding of extinction learning by demonstrating that there is individual variation in extinction and relapse processes and highlight potential difficulties in applying extinction-based therapies to drug addiction treatment in the clinic.

## 1. Introduction

The attribution of incentive-motivational value to drug-related cues is believed to underlie craving and relapse in addiction (Robinson and Berridge, 1993). In support of this, reactivity to drug-related cues before and during treatment is associated with craving (Witteman et al., 2015) and relapse (Garland et al., 2012; Janes et al., 2010; Papachristou et al., 2014). Reducing the strength of drug-related memories, for example through extinction learning, is one way to decrease craving and facilitate addiction treatment. In the context of addiction, extinction is defined as a learning process that reduces the frequency or intensity of conditioned responses (CRs; e.g., craving) to drug-related cues. During extinction learning, the drug-related cue, or conditioned stimulus (CS; e.g., drug paraphernalia), is presented in the absence of the drug, or unconditioned stimulus (US; e.g., cocaine), which results in a new learning process that inhibits expression of the original learning (Bouton, 2004; Delamater, 2004; Myers and Davis, 2002).

Because patients suffering from addiction exhibit enduring cue reactivity even after treatment, a better understanding of extinction learning is required to increase its effectiveness in therapeutic settings. In the 1970s, individual variation was observed in abstinence patterns following behavioral treatment for addiction, and it was hypothesized that relapse was causally related to inadequate extinction learning established during treatment (Hunt et al., 1971; Litman et al., 1979). During the 1990s, cue-exposure therapy was advocated as a potential psychotherapy for addiction (Heather and Bradley, 1990), and initial studies appeared promising in decreasing cue reactivity (Franken et al., 1999) and relapse (Drummond and Glautier, 1994). Subsequent studies, however, did not replicate these findings, and a meta-analysis of cue-exposure studies showed no consistent evidence of efficacy (Conklin and Tiffany, 2002). However, animal extinction research upon which these treatments were based has advanced tremendously since that time, and exposure-based therapies might be informed and improved by novel discoveries in extinction learning. For example, little is known regarding how reward-related cues that are imbued with incentive-motivational value extinguish and relapse.

Pavlovian conditioned approach (PCA) procedures can be used to investigate individual variation in the attribution of incentive-motivational value to reward-related cues (Robinson and Flagel, 2009). During PCA training, a CS (e.g., a lever) response-independently predicts the delivery of an US (e.g., a food pellet). During training, three patterns of CRs develop: sign-tracking (CS-directed CRs), goal-tracking (US-directed CRs), and an intermediate response (both CRs). Previous research has shown that sign-trackers (STs), but not goal-trackers (GTs), attribute reward-related cues with incentive-motivational value, transforming them into powerful motivators of behavior (Flagel et al., 2009) that can promote addiction-like behaviors, such as cue-induced reinstatement of drug-seeking (Saunders and Robinson, 2010; Yager et al., 2015). It is also clear that sign-tracking behaviors directed toward reward-related cues imbued with incentive-motivational value are more resistant to extinction than goal-tracking behavior (Ahrens et al., 2016; Beckmann and Chow, 2015). It is unknown, however, (1) how long resistance to extinction of sign-tracking behavior persists, (2) whether PCA phenotypes differentially undergo relapse processes like reinstatement (recovery of conditioned responding as a result of US presentation) and spontaneous recovery (recovery of conditioned responding due to the passage of time), or (3) whether drugs currently being investigated as cognitive enhancers during extinction-based therapies are effective in promoting extinction of sign-tracking behavior (Quirk and Mueller, 2008).

To address this, rats received PCA training followed by extinction training and two tests for relapse-related processes: reinstatement and spontaneous recovery. In addition, a separate group of rats underwent PCA training followed by operant conditioning, extinction and tests for reinstatement and spontaneous recovery. Operant procedures were performed in parallel to determine whether differences in extinction learning are unique to Pavlovian procedures or a generalized feature of associative learning. Finally, three cognitive enhancers—sodium butyrate (NaB; a histone deacetylase inhibitor), D-cycloserine (DCS; an N-methyl-D-aspartate receptor [NMDAR] partial agonist), and fibroblast growth factor 2 (FGF2)—were administered during extinction training to facilitate the extinction of sign-tracking behavior. These drugs were selected because they have been previously shown to enhance extinction learning in animal models (Graham and Richardson, 2009; 2010; Paolone et al., 2009; Zhu et al., 2017).

## 2. Methods and Materials

### 2.1. Animals

One hundred sixty-eight adult male Sprague Dawley rats (275-300 g) were purchased from Charles River Laboratories and Envigo (formerly known as Harlan Laboratories). In Experiment 1, rats were used equally from Charles River Laboratories (Barriers C72, R04, and R09) and Envigo (202A, 202C, 217, and 206). In Experiment 2, rats were used exclusively from Charles River Laboratories (Barrier C72). Rats were maintained on a 12 h light/dark cycle, and standard rodent chow and water were available *ad libitum.* All procedures were approved by the University Committee on the Use and Care of Animals (University of Michigan; Ann Arbor, MI).

### 2.2. Drugs

Sodium butyrate (NaB; #sc-202341; Santa Cruz Biotechnology, Inc.; Dallas, TX), D-cycloserine (DCS; #C6880; Sigma-Aldrich, Inc.; St. Louis, MO), and fibroblast growth factor 2 (FGF2; #45103P; QED Bioscience, Inc.; San Diego, CA) were used. In Experiment 1, NaB (200 mg/kg, i.p.; 10 mL/kg) was dissolved in saline (pH = 7.34), which was also used as the vehicle control. In Experiment 2, DCS (30 mg/kg, i.p.; 1 mL/kg) was dissolved in saline (pH = 7.34) and FGF2 (20 μg/kg, i.p.; 2 mL/kg) was dissolved in 0.1% bovine serum albumin (Sigma Aldrich, Inc.) in 0.1 M phosphate-buffered saline (pH = 7.34). A combination of saline (1 mL/kg, i.p.) and 0.1% bovine serum albumin in 0.1 M phosphate-buffered saline (1 mL/kg, s.c.) was used as the vehicle control, and drug groups received drug and the opposite vehicle, so that each group had the same amount of injections. Doses of DCS (30 mg/kg; i.p.) and FGF2 (20 μg/kg; s.c.) were selected based upon previous studies showing that these doses enhance extinction training (Graham and Richardson, 2009; Vurbic et al., 2011). The dose of NaB (200 mg/kg; i.p.) was selected because it has been shown to enhance experience-dependent histone acetylation (Kumar et al., 2005).

### 2.3. Apparatus

Modular conditioning chambers (24.1 cm width × 20.5 cm depth × 29.2 cm height; MED Associates, Inc.; St. Albans, VT) were used for Pavlovian and operant conditioning. Each chamber was in a sound-attenuating cubicle equipped with a ventilation fan to provide ambient white noise. For Pavlovian conditioning, chambers were equipped with a pellet magazine, an illuminated retractable lever (counterbalanced on the left or right of the pellet magazine), and a red house light on the wall opposite to the pellet magazine. When inserted into the chamber, the retractable lever was illuminated by an LED light within the lever housing. A pellet dispenser delivered banana-flavored food pellets into the pellet magazine, and an infrared sensor inside the pellet magazine detected head entries. For operant conditioning, the lever was removed from the chamber and replaced with two nose-poke ports on either side of the pellet magazine.

### 2.4. Pavlovian Conditioned Approach: Procedure

For two days prior to pretraining, rats were familiarized with banana-flavored food pellets (45 mg; Bioserv; Frenchtown, NJ) in their home cages. Twenty-four hours later, rats were placed into the operant chambers and underwent one pretraining session during which the red house-light remained on, but the lever was retracted. Fifty food pellets were delivered on a variable time (VT) 30 schedule (i.e., one food pellet was delivered on average every 30 s, but actual delivery varied between 0-60 s). All rats consumed all the food pellets by the end of the pretraining session. Twenty-four hours later, rats underwent daily PCA training sessions over seven days. Each trial during a test session consisted of extension of the illuminated lever (the CS) into the chamber for 8 s on a VT 90 schedule (30-150 s). Retraction of the lever was immediately followed by the response-independent delivery of one food pellet (the US) into the pellet magazine. Each test session consisted of 25 trials of CS-US pairings, resulting in a total session length of approximately 40 min. All rats consumed all the food pellets that were delivered.

Following PCA training, approximately half of the rats underwent 29 daily PCA extinction sessions during which the CS was presented but not did result in the delivery of a food pellet. Next, rats underwent a reinstatement test during which five food pellets were delivered at the beginning of the session followed immediately by an extinction session. Rats then underwent 10 more PCA extinction sessions followed by a spontaneous recovery test two weeks later. During the spontaneous recovery test, rats were placed back into chambers and behavior was recorded under extinction conditions. During Pavlovian extinction, reinstatement, and spontaneous recovery, the first three CRs were counted for statistical analysis. Probability of CRs was used as the dependent measure so that sign- and goal-tracking could be directly compared. For detailed analysis of extinction recall, the first and last CRs of sessions were counted for statistical analysis, and number of CRs was used as the dependent measure.

### 2.5. Operant Conditioning: Procedure

Following PCA training, approximately half of the rats underwent 12 operant conditioning sessions. Chocolate-flavored food pellets were used instead of banana-flavored pellets so that Pavlovian and operant procedures would not have the same US. Operant conditioning consisted of a 50-min session with three within-session phases: food pellet delivery on a fixed ratio (FR) 1 schedule for the first five food pellet deliveries (i.e., one active nose-poke response resulted in food pellet delivery), an FR 7 schedule for the next five food pellet deliveries (i.e., seven active nose-poke responses resulted in pellet delivery), then a variable ratio 20 (VR 20) schedule for all further food pellet deliveries (i.e., the number of active nose-poke responses required to result in pellet delivery varied randomly between 5 and 35). Following operant conditioning, rats underwent six sessions of extinction training (50 min) during which active nose-poke responses no longer resulted in food pellet delivery. Next, rats underwent a reinstatement test during which five food pellets were delivered at the beginning of the session followed immediately by an extinction session. Rats then underwent two more operant extinction sessions followed by a spontaneous recovery test two weeks later. During the spontaneous recovery test, rats were placed back into chambers, and behavior was recorded under extinction conditions.

### 2.6. Statistical Analysis

PCA behavior was scored using an index that combines the number, latency, and probability of lever presses (sign-tracking CRs) and magazine entries (goal-tracking CRs) during CS presentations within a session. Briefly, we averaged together the response bias (i.e., number of lever presses and magazine entries for a session; [lever presses – magazine entries] / [lever presses + magazine entries]), latency score (i.e., average latency to perform a lever press or magazine entry during a session; [magazine entry latency – lever press latency]/8 s), and probability difference (i.e., proportion of lever presses or magazine entries; lever press probability – magazine entry probability). The index scores behavior from +1.0 (absolute sign-tracking) to −1.0 (absolute goal-tracking), with 0 representing no bias (Meyer et al., 2012). Rats were classified using the following cut-offs: STs (x ≥ 0.5), IRs (−0.5 < × < 0.5), and GTs (x ≤ – 0.5).

SPSS (Version 22; IBM, Inc.) was used for all statistical analysis. For all linear mixed models, the covariance structure was selected based upon Akaike’s information criterion (i.e., the lowest number criterion represents the highest quality statistical model using a given covariance structure). For PCA and operant training and extinction, behavior across training sessions was analyzed using a linear mixed model with an autoregressive (AR1) covariance structure. For Pavlovian extinction recall as well as reinstatement and spontaneous recovery tests, two-way analysis of variance (ANOVA) was performed. With a significant ANOVA, multiple comparisons were performed using Fisher’s Least Significant Difference (LSD) post hoc test.

## 3. Results

### 3.1. Experiment 1: PCA training

Rats underwent seven daily sessions of PCA training and were classified as STs (n = 36), intermediate responders (IRs; n = 29), and GTs (n = 30) based upon their average PCA index scores over Sessions 6 and 7. Figure 1 shows that there was an effect of Phenotype on lever press number (F_(2,130.74)_ = 95.2, p = 3.0 × 10^-26^), latency (F_(2,114.31)_ = 38.4, p = 1.8 × 10^-13^), and probability (F_(2,124)_ = 101, p = 1.1 × 10^-26^) as well as magazine entry number (F_(2,123.63)_ = 51.3, p = 6.2 × 10^-17^), latency (F_(2,127.70)_ = 70.9, p = 1.9 × 10^-21^), and probability (F_(2,124.89)_ = 81.7, p = 2.1 × 10^-23^). Following PCA training, rats were divided into two groups. The first group underwent PCA extinction and testing for reinstatement and spontaneous recovery (ST, n = 18; IR, n = 17, GT, n = 12); the second group underwent operant training, extinction, and testing for reinstatement and spontaneous recovery (ST, n = 18; IR, n = 12, GT, n = 18).

**Figure 1.**
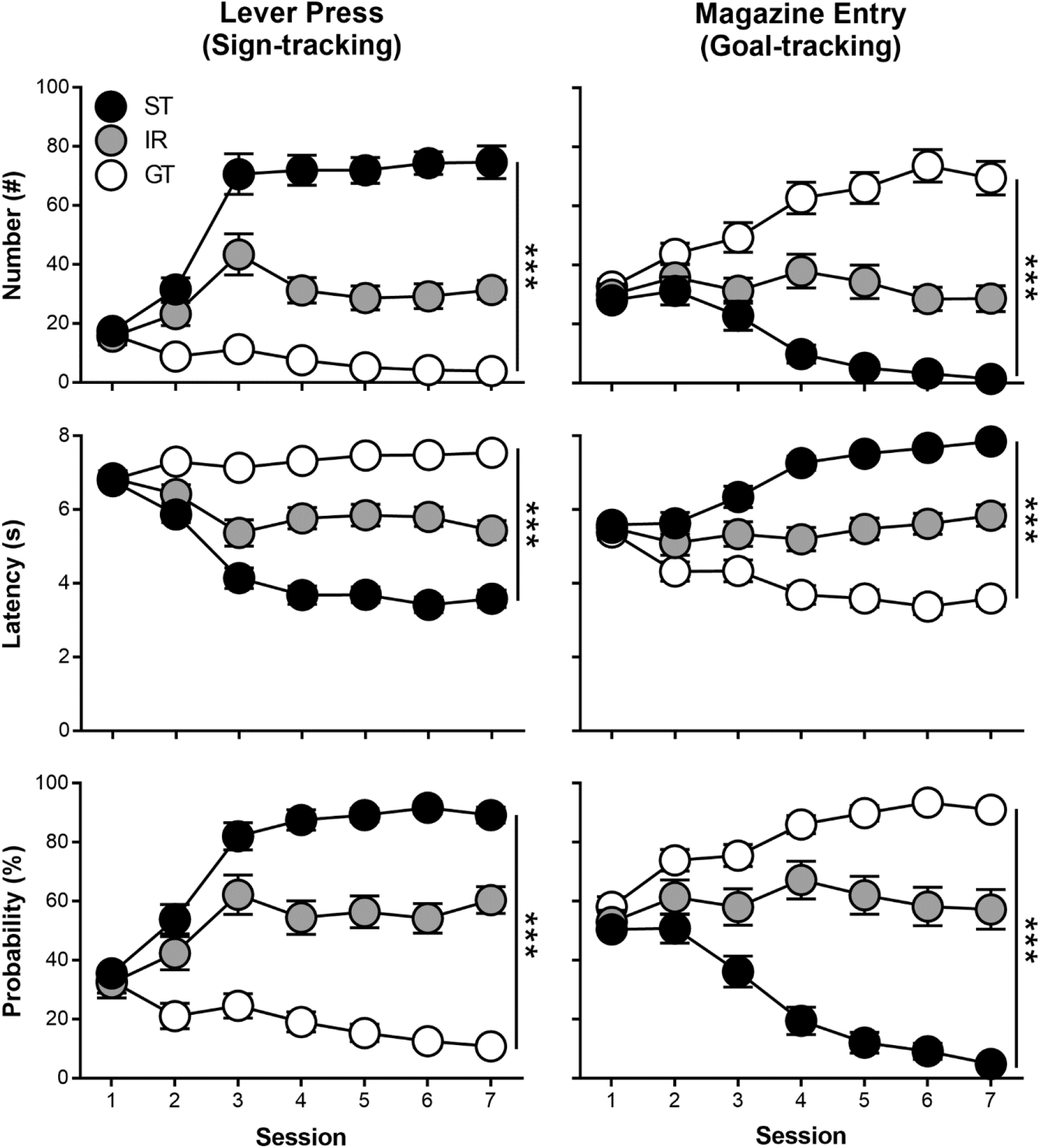
Rats underwent Pavlovian conditioned approach training over seven daily sessions and were classified as sign-trackers (STs), intermediate-responders (IRs), or goal-trackers (GTs) based on the average lever press and magazine entry number, latency, and probability during Sessions 6 and 7. Data are presented as mean and S.E.M. *** – p < 0.001.

### 3.2. Experiment 1: PCA extinction, reinstatement, and spontaneous recovery

Figure 2 shows extinction and extinction recall of PCA behavior in STs (n = 18), IRs (n = 17), and GTs (n = 12). Over 29 daily sessions of PCA extinction, STs and GTs extinguished their conditioned responding (Figure 2A; effect of Session: F_(28,407.14)_ = 10.1, p = 5.3 × 10^-32^); however, extinction of sign-tracking CRs in STs was weaker than that of goal-tracking CRs in GTs (effect of Phenotype: F_(1,223.78)_ = 55.5, p = 2.0 × 10^-12^). In addition, IRs extinguished their conditioned responding (Figure 2B; effect of Session: F_(28,497.57)_ = 5.13, p = 5.6 × 10^-15^), but extinction of sign-tracking CRs was weaker than that of goal-tracking CRs in IRs (effect of Response: F_(1,184.56)_ = 89.5, p = 1.5 × 10^-17^). Analysis of the first and last CS trials during the last session of PCA training and the first four sessions of PCA extinction revealed that STs extinguished sign-tracking within sessions, but did not extinguish it between sessions (Figure 2C; effect of CS: F_(1,170)_ = 132.27, p = 5.21 × 10^-23^). Post hoc comparisons revealed that STs sign-tracked less on the last CS trial compared to the first CS trial on every extinction session (ps < 0.05); however, sign-tracking on the first CS trial during extinction sessions was not different from the last PCA session (ps > 0.05). Similarly, IRs extinguished sign-tracking within-session, but did not extinguish it between-session (Figure 2E; effect of CS: F_(1,110)_ = 24.90, p = 4.08 × 10^-11^). Post hoc comparisons revealed that IRs sign-tracked less on the last CS trial compared to the first CS trial on every extinction session (ps < 0.05), but sign-tracking on the first CS trial during extinction sessions was not different from the last PCA session (ps > 0.05). In contrast, GTs extinguished goal-tracking both within sessions and between sessions (Figure 2D; effect of CS: F_(1,110)_ = 11. 14, p = 0.001). Post hoc comparisons revealed that GTs goal-tracked less on the last CS trial compared to the first CS trial on the first extinction session (p < 0.001), and goal-tracking on subsequent sessions was low and similar between the first and last CS trials (ps > 0.05). In addition, goal-tracking on the first CS trial of Extinction Sessions 2-4 were significantly lower than the first CS trial of the last PCA session (ps < 0.01). Finally, IRs extinguished their goal-tracking within sessions, but not between sessions (Figure 2E; effect of CS: F_(1,50)_ = 14.25, p = 2.21 × 10^-4^). Pos hoc comparisons revealed that IRs goal-tracked less on the last CS trial compared to the first CS trial on the first extinction session (p < 0.01), and goal-tracking on subsequent sessions was low and similar between the first and last CS trials (ps > 0.05). Goal-tracking on the first CS trial of the last PCA session and extinction sessions, however, was too low to observe between-session extinction (ps > 0.05).

**Figure 2.**
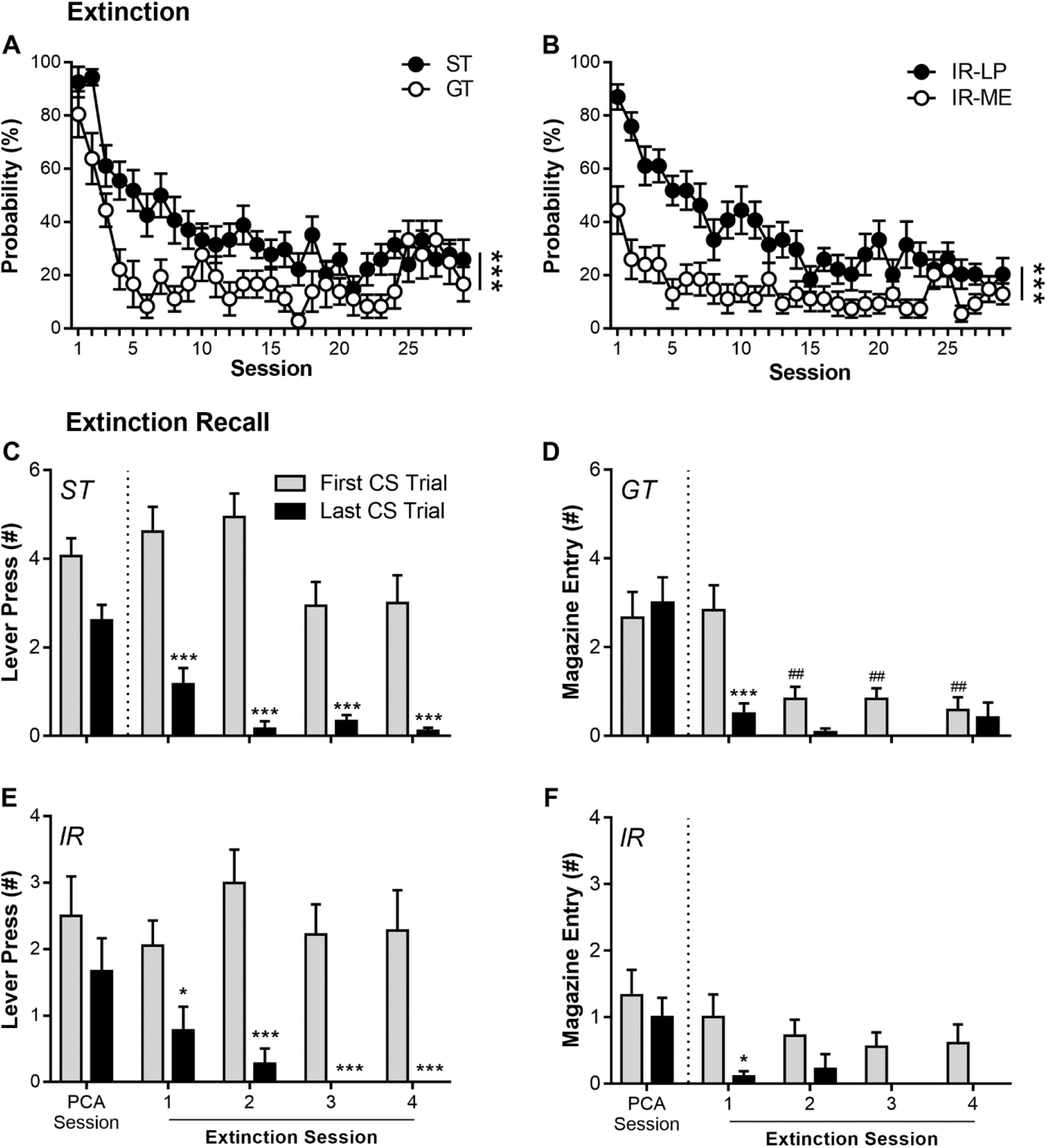
Sign-trackers (STs), intermediate-responders (IRs), and goal-trackers (GTs) were tested for extinction of Pavlovian conditioned approach (PCA) behavior. (A-B) Rats underwent 29 PCA extinction sessions, and the probabilities of sign-tracking conditioned responses (lever press; LP) in STs, goal-tracking conditioned responses (magazine entry; ME) in GTs, and both conditioned responses in IRs were compared. (C-F) To investigate extinction recall of PCA behavior, the first and last conditioned stimulus (CS) within a session were compared between the last session of PCA training and the first four sessions of PCA extinction. Data are presented as mean and S.E.M. * – p < 0.05, *** – p < 0.001, within-subjects comparisons; ## – p < 0.01, between-subjects comparison.

Figure 3 shows tests of reinstatement and spontaneous recovery following PCA extinction sessions. Behavior was compared against the last PCA extinction session (baseline). For GTs and STs, exposure to the US did not reinstate behavior (Figure 3A; effect of Reinstatement: F_(1,53)_ = 3.90, p = 0.054; effect of Phenotype: F_(1,53)_ = 3.57, p = 0.064; interaction of Phenotype x Reinstatement: F_(1,53)_ = 0.004, p = 0.95). For IRs, exposure to the US reinstated behavior (Figure 3B; effect of Reinstatement: F_(1,63)_ = 15.1, p = 2.5 × 10^-4^), and post hoc comparisons revealed that sign-tracking CRs (p = 0.013) and goal-tracking CRs (p = 0.005) both reinstated. There were no differences, however, between the reinstatement of sign- and goal-tracking CRs (effect of Phenotype: F_(1,63)_ = 0.69, p = 0.41; interaction of Phenotype x Reinstatement: F_(1,63)_ = 0.077, p = 0.78). Following the reinstatement test, STs and GTs extinguished conditioned responding over 10 daily PCA extinction training sessions (data not shown; effect of Session: F_(9,165.23)_ = 2.54, p = 0.009), and STs continued to extinguish their lever-press CRs slower than GTs extinguished their magazine-entry CRs (effect of Phenotype: F_(1,73.87)_ = 13.2, p = 5.3 × 10^-4^; interaction of Phenotype x Session: F_(1,165.24)_ = 0.45, p = 0.91). Moreover, IRs extinguished conditioned responding as well (effect of Session: F_(1,192.47)_ = 1.97, p = 0.044), and they continued to extinguish their lever-press CRs slower than their magazine-entry CRs (effect of Response: F_(1,70.16)_ = 13.8, p = 4.1 × 10^-4^; interaction of Response x Session: F_(1,192.47)_ = 0.59, p = 0.80).

**Figure 3.**
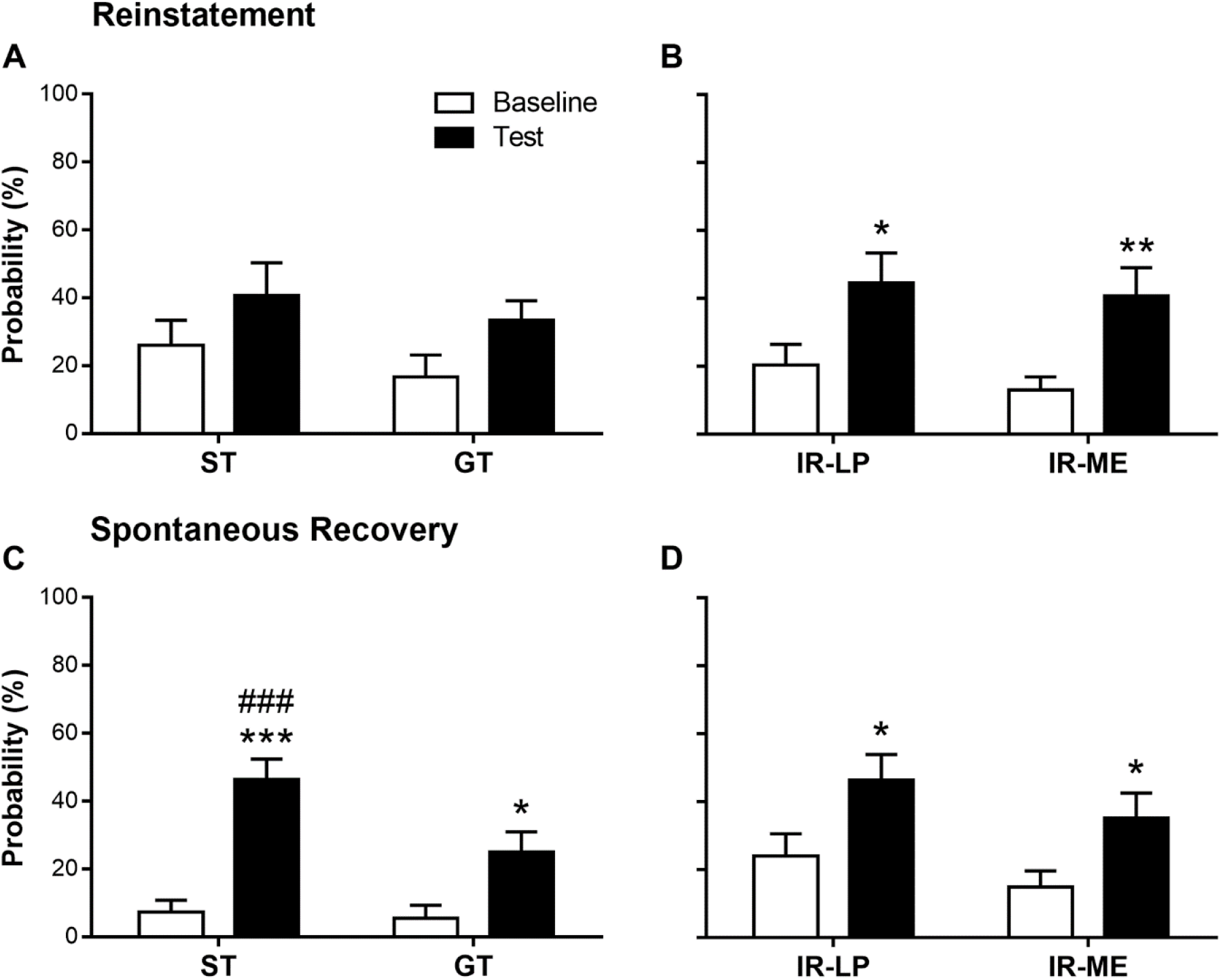
Following extinction of Pavlovian conditioned approach behavior, sign-trackers (STs), intermediate-responders (IRs) and goal-trackers (GTs) underwent tests for (C-D) reinstatement and (E-F) spontaneous recovery of lever press (LP) and magazine entry (ME) conditioned responses. Baseline refers to the most recent PCA extinction training session relative to the specific test session. Data are presented as mean and S.E.M. * – p < 0.05, ** – p < 0.01, *** – p < 0.001, within-subjects comparisons; ### – p < 0.001, between-subjects comparison.

Two weeks after PCA re-extinction training, rats underwent a spontaneous recovery test, and behavior was compared against the last PCA extinction session (baseline). For GTs and STs, conditioned responding spontaneously recovered (Figure 3C; effect of Session: F_(1,53)_ = 36.3, p = 1.7 × 10^-7^), but CRs recovered differently (effect of Phenotype: F_(1,53)_ = 6.87, p = 0.011; interaction of Phenotype x Session: F_(1,53)_ = 4.66, p = 0.035). Post hoc comparisons revealed that STs (p = 4.8 × 10^-8^) and GTs (p = 0.015) both spontaneously recovered their respective CRs; however, STs spontaneously recovered conditioned responding to a greater degree (p = 0.0013). For IRs, conditioned responding spontaneously recovered (Figure 3D; effect of Session; F_(1,63)_ = 10.8, p = 0.002), and post hoc comparisons revealed that sign-tracking CRs (p = 0.018) and goal-tracking CRs (p = 0.03) both recovered. There were no differences, however, between the spontaneous recovery of sign- and goal-tracking CRs in the IRs (effect of Phenotype: F_(1,63)_ = 2.48, p = 0.12; interaction of Phenotype x Reinstatement: F_(1,63)_ = 0.02, p = 0.89).

### 3.3. Experiment 1: Operant conditioning, extinction, reinstatement, and spontaneous recovery

Figure 4*A* shows that over 12 sessions of operant conditioning rats (STs, n = 18; IRs, n = 12; GTs, n = 18) successfully acquired the operant response (effect of Session: F_(11, 427.97)_ = 15.1, p = 8.3 × 10^-25^) and discriminated between active and inactive nose-poke ports (effect of Port: F_(1, 129.97)_ = 219, p = 1.2 × 10^-24^). In addition, there were no differences in active nose-poke responses between phenotypes (effect of Phenotype: F_(2,129.97)_ = 1.98, p = 0.14). Following operant training, rats underwent six operant extinction sessions, during which rats successfully extinguished their active nose-poke responses (Figure *4B*: effect of Session; F_(5, 298.32)_ = 66.4, p = 1.9 × 10^-46^). There was no difference in extinction between phenotypes (effect of Phenotype: F_(2, 71.74)_ = 0.34, p = 0.72). Following operant extinction, rats underwent a reinstatement test, and their behavior was compared against the last operant extinction training session (baseline). Exposure to the US reinstated behavior (Figure 4C: effect of Session; F_(1,90)_ = 8.48, p = 0.0045); however, post hoc comparisons revealed that only GTs (p = 0.012), but not IRs (p = 0.11) or STs (p = 0.37), increased their active nose-poke responses. Following the reinstatement test, rats were extinguished over two daily operant extinction training sessions. After two weeks following operant extinction training, rats underwent a spontaneous recovery test, and behavior was compared against the last operant extinction training session (baseline). Operant responding spontaneously recovered (Figure 4D: effect of Session; F_(1,90)_ = 10.2, p = 0.002), and post hoc comparisons revealed that only GTs (p = 0.023), but not IRs (p = 0.09) or STs (p = 0.12), increased their active nose-poke responses.

**Figure 4.**
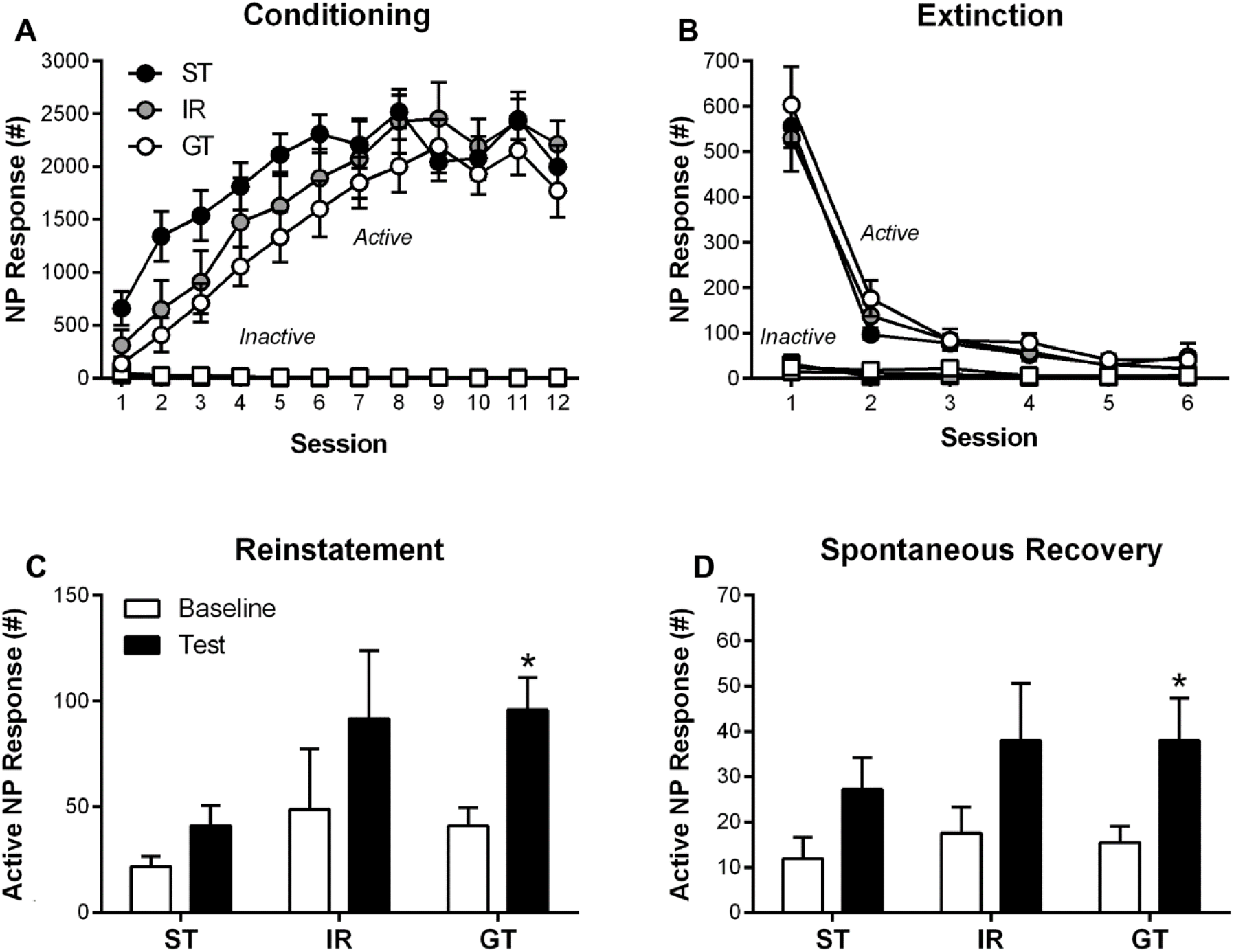
Sign-trackers (STs), intermediate-responders (IRs), and goal-trackers (GTs) were tested for operant conditioning, extinction, reinstatement, and spontaneous recovery, and their active and inactive nose-poke (NP) responses were recorded during each session. Baseline refers to the most recent operant extinction training session relative to the specific test session. Data are presented as mean and S.E.M. * – p < 0.05.

### 3.4. Experiment 2: The effect of sodium butyrate on PCA extinction

Seven days following the spontaneous recovery test in Experiment 1, rats underwent three daily additional PCA training sessions to restore conditioned responding followed by four daily PCA extinctions. After each PCA extinction session, rats immediately received NaB or saline. Figure *5A-B* shows that STs and GTs extinguished their conditioned responding (effect of Session: F_(3,109.67)_ = 6.50, p = 4.4 × 10^-4^), but sign-tracking CRs in STs were more resistant to extinction than goal-tracking CRs in GTs (effect of Phenotype: F_(1, 59.57)_ = 17.6, p = 1.1 × 10^-4^). However, NaB did not affect extinction training in either STs or GTs (effect of Drug: F_(2, 49.45)_ = 0.16, p = 0.90; interaction of Drug x Phenotype: F_(1, 49.47)_ = 0.16, p = 0.69; interaction of Drug x Phenotype x Session: F_(6, 109.67)_ = 0.61, p = 0.61). Similarly, IRs extinguished their conditioned responding (effect of Session: F_(3,95.62)_ = 6.40, p = 0.001), and sign-tracking CRs were more resistant to extinction than goal-tracking CRs in IRs (effect of Response: F_(1,40.85)_ = 30.0, p = 2.4 × 10^-6^). However, NaB did not affect extinction training in IRs (effect of Drug: F_(1, 40.86)_ = 3.02, p = 0.09; interaction of Drug x Response: F_(1, 40.85)_ = 0.095, p = 0.76; interaction of Drug x Response x Session: F_(3, 95.62)_ = 0.93, p = 0.43). Also, differences in prior behavioral training and duration of testing between the operant and Pavlovian training groups did not affect the conditioned responding during extinction training of sign-tracking in STs and goal-tracking in GTs (effect of Cohort: F_(1,49.45)_ = 0.74, p = 0.39) or sign- and goal-tracking in IRs (effect of Cohort: F_(1,34.02)_ = 0.67, p = 0.42).

**Figure 5.**
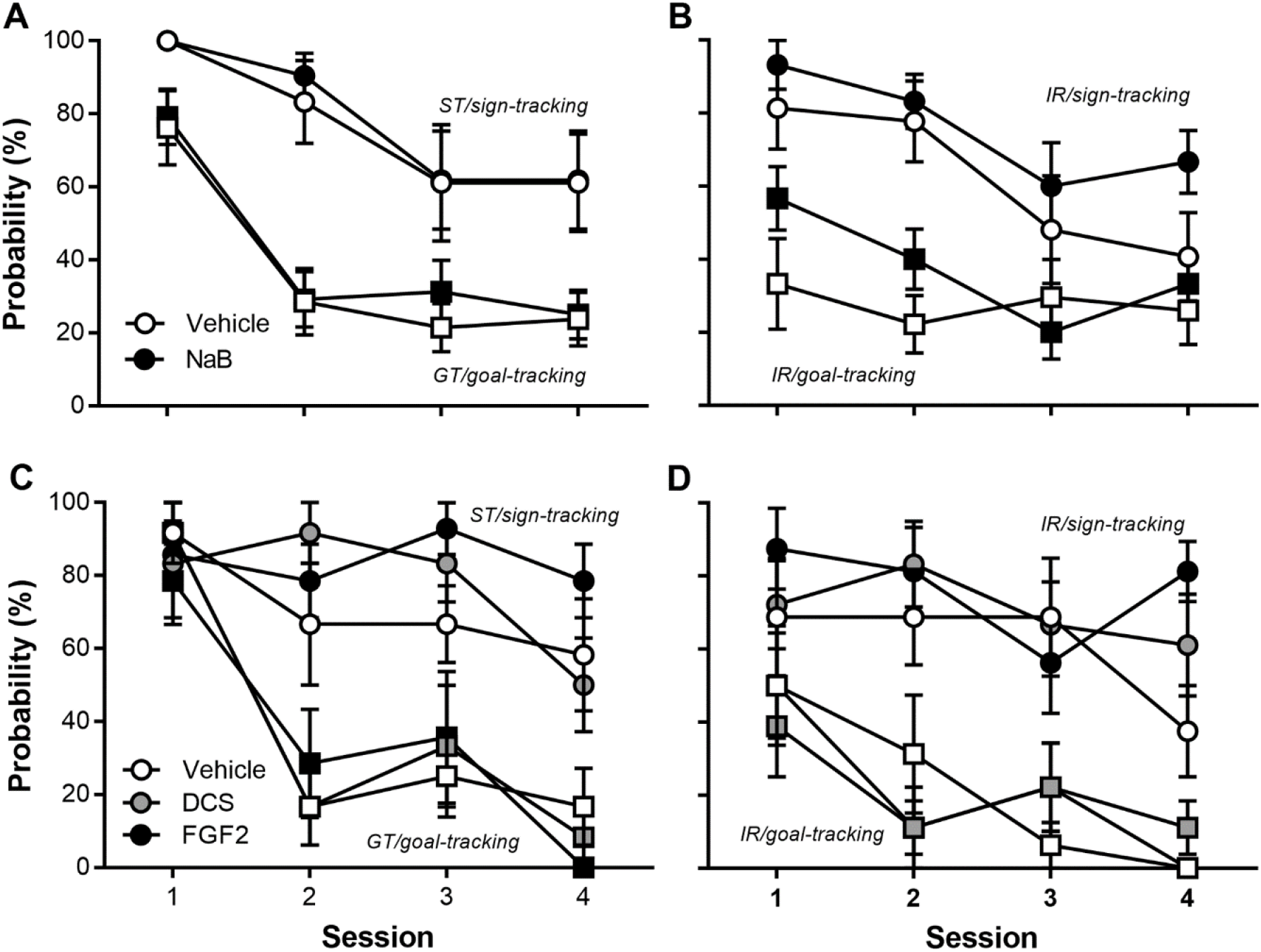
Sign-trackers (STs), goal-trackers (GTs), and intermediate-responders (IRs) underwent four sessions of Pavlovian extinction. Immediately following each session, rats were administered (A-B) sodium butyrate (NaB; 200 mg/kg, i.p.), (C-D) D-cycloserine (DCS; 30 mg/kg, i.p.) or fibroblast growth factor 2 (FGF2; 20 μg/kg, s.c.), and their respective vehicles. Data are presented as mean and S.E.M.

### 3.5. Experiment 2: The effect of DCS and FGF2 on PCA extinction

In a new cohort, rats underwent seven daily sessions of PCA training and were classified as STs (n = 19), IRs (n = 26), and GTs (n = 19) based upon their average PCA index scores over Sessions 6 and 7. There was an effect of Phenotype on lever press number (F_(2,85.77)_ = 52.8, p = 1.14 × 10^-15^), latency (F_(2,86.35)_ = 71.1, p = 5.58 × 10^-19^), and probability (F_(2,84.56)_ = 88.9, p = 1.6 × 10^-21^) as well as magazine entry number (F_(2,118.64)_ = 38.0, p = 1.74 × 10^-13^), latency (F_(2,88.59)_ = 42.0, p = 1.43 × 10^-13^), and probability (F_(2,117.19)_ = 61.0, p = 6.99 × 10^-19^). Twenty-four hours following PCA training, rats were divided into two groups and underwent four daily sessions of PCA extinction, each followed immediately by a vehicle, DCS, or FGF2 injection. As mentioned previously, because DCS and FGF2 have different vehicle solutions, vehicle-treated rats received two vehicle injections and drug-treated rats received drug and the additional vehicle solution. Figure *5C-D* shows that STs and GTs extinguished their conditioned responding (effect of Session: F_(3,93.16)_ = 24.3, p = 1.07 × 10^-11^), but sign-tracking CRs in STs were more resistant to extinction than goal-tracking CRs in GTs (effect of Phenotype: F_(1,44.05)_ = 27.7, p = 4.56 × 10^-8^). In addition, DCS or FGF2 did not affect extinction training in either STs or GTs (effect of Drug: F_(2,44.05)_ = 0.73, p = 0.49; interaction of Drug x Phenotype: F_(1,44.05)_ = 0.98, p = 0.38; interaction of Drug x Phenotype x Session: F_(6,93.16)_ = 0.77, p = 0.60). Similarly, IRs extinguished their conditioned responding (effect of Session: F_(3,130.42)_ = 8.39, p = 3.85 × 10^-5^), and sign-tracking CRs were more resistant to extinction than goal-tracking CRs in IRs (effect of Response: F_(1,60.53)_ = 79.9, p = 1.16 × 10^-12^). In addition, DCS or FGF2 did not affect extinction training of either CR in IRs (effect of Drug; F_(2, 60.53)_ = 0.31, p = 0.73: interaction of Drug x Response: F_(1, 60.53)_ = 0.66, p = 0.52; interaction of Drug x Response x Session: F_(6,130.42)_ = 0.48, p = 0.82).

## 4. Discussion

In the current study, we demonstrated individual differences in Pavlovian extinction and spontaneous recovery of reward-related cues that have or have not been imbued with incentive-motivational value. For example, sign-tracking behavior in STs was more resistant to extinction than goal-tracking in GTs. Similarly, sign-tracking behavior in IRs was more resistant to extinction than goal-tracking behavior. Furthermore, these extinction deficits in sign-tracking were the result of between-session deficits in extinction recall whereas within-session extinction was relatively unimpaired. In addition, sign-tracking behavior in STs exhibited higher spontaneous recovery than goal-tracking behavior in GTs following extinction. In contrast, operant conditioning and extinction were not different between phenotypes, which supports previous findings that they do not differ in operant learning (Saunders and Robinson, 2010; 2011). Thus, extinction deficits in STs are specific to Pavlovian procedures during which reward-related cues are imbued with incentive-motivational value (Ahrens et al., 2016; Beckmann and Chow, 2015).

Significant individual differences in extinction learning have been observed in rodents (Shumake et al., 2014) and humans (Gershman and Hartley, 2015; Lommen et al., 2013). Contemporary learning theory posits that extinction learning likely involves the formation of new CS-no US associations that inhibit the original CS-US associations (Konorski, 1948; Rescorla, 1993; Rescorla, 2001; Hall, 2002). One understudied aspect of extinction, however, is whether it uniformly influences different associations between the CS and US (i.e., sensory and motivational properties; Konorski, 1967; Delamater and Oakeshott, 2007), which would have important implications for learning theory and the study of individual differences. Certain theories of extinction (i.e., AESOP theory) have acknowledged the utility of understanding these differences (Wagner and Brandon, 1989), and PCA procedures can separate at least two associations between the CS and US: incentive-motivational (sign-tracking CR) and sensory-predictive (goal-tracking CR).

In the present study, the extinction of goal-tracking generally supports the Rescorla-Wagner model (Rescorla and Wagner, 1972) model and negative prediction errors (Delamater and Westbrook, 2014; Schultz, 1998; Schultz et al., 1997). The extinction of sign-tracking, however, is not as consistent with the assumptions of the Rescorla-Wagner model (i.e., extinction results in decrements in associative strength caused by negative prediction errors). Although it is currently unknown what behavioral and neurobiological mechanisms underlie the extinction of sign-tracking, our results may point toward deficient memory consolidation, because sign-tracking extinguishes within but not between sessions. One possibility is that the attribution of incentive-motivational value to a CS impairs extinction learning by increasing attention to the CS (Robbins, 1990).

There are several brain regions that could be responsible for resistance to extinction of sign-tracking behavior. First, it may result from hypoactivity in the prefrontal cortex (PFC), because it has previously been shown that temporary inactivation or NMDAR antagonism of the nidopallium caudolaterale (the functional analogue of the mammalian PFC) impairs the extinction of sign-tracking behavior in pigeons (Lengersdorf et al., 2015; Lengersdorf et al., 2014). Within-session extinction does not require the infralimbic (IL) cortex of the PFC; however, consolidation of within-session extinction does require the IL (Quirk et al., 2000). Because sign-tracking rats have decreased levels of BDNF in the PFC (Morrow et al., 2015), and BDNF in the IL is both necessary and sufficient for Pavlovian extinction (Rosas-Vidal et al., 2014), it is possible that decreased levels of BDNF in the IL underlie extinction deficits in STs. Second, resistance to extinction could result from persistent cue-induced activation of the basolateral amygdala (BLA), which has increased neural activity in STs relative to GTs under acute extinction conditions (Yager et al., 2015). Alternatively, abnormal activity during extinction learning of sign-tracking behavior may occur in both regions, because it is known that a BLA-IL circuit contributes to the formation and extinction of reward-related memories (Rosen et al., 2015; Xin et al., 2014). Interestingly, IRs showed similar resistance to extinction of sign-but not goal-tracking behavior, suggesting that these circuits may be differentially engaged between the two behaviors in IRs and supporting the notion that sign- and goal-tracking behaviors are learned and extinguished through unique and dissociable systems even in the same animal (Clark et al., 2012).

As previously mentioned, increased spontaneous recovery of sign-tracking in STs compared to goal-tracking in GTs may result from the same processes that cause individual variation in resistance to extinction. Spontaneous recovery results from the fact that extinction memories inhibit, rather than erase, conditioning memories and are susceptible to weakening over time (Todd et al., 2014). Previously, it has been demonstrated that extinction is associated with increased excitability of IL neurons, which is reversed during spontaneous recovery (Cruz et al., 2014); therefore, resistance to extinction and enhanced spontaneous recovery may stem from similar impairments in synaptic plasticity in the IL. Moreover, and as previously described, STs have increased neural activity in the BLA relative to GTs under acute extinction conditions, and BLA activity during extinction predicts the strength of subsequent spontaneous recovery (Courtin et al., 2014) whereas its inactivation decreases spontaneous recovery (Peters et al., 2008).

Recently, there has been a great interest in using cognitive enhancers to facilitate extinction learning (Kaplan and Moore, 2011). In the current study, three promising cognitive enhancers were given during the extinction of sign- and goal-tracking behaviors, yet none of these compounds facilitated extinction learning under these testing conditions. Interestingly, all three were unsuccessful even though they enhance extinction learning through separate mechanisms. Each compound either increases synaptic plasticity through glutamate receptor modulation (DCS), histone acetylation (NaB), or growth factor-mediated intracellular signaling (FGF2).

There are several important considerations as to why these cognitive enhancers were ineffective at promoting extinction of sign-tracking. First, the vast majority of research investigating Pavlovian extinction learning and its pharmacological enhancement have stemmed from fear conditioning in animals (Todd et al., 2014) and anxiety disorders in humans (Hofmann et al., 2015). Although it is known that there is an overlap in the circuits (Peters et al., 2009) and signaling pathways (Heldt et al., 2014) underlying appetitive and aversive learning, it is also known that there are important differences in these processes (Hebart and Glascher, 2015; Knapska et al., 2006; Reichelt and Lee, 2013). Indeed, most behavioral studies of extinction stem from appetitive conditioning procedures whereas most neuropharmacological studies of extinction stem from aversive conditioning procedures (Delamater, 2004). Second, there is a possibility that these drugs might have been effective if given before Pavlovian extinction training. In the current study, we targeted the consolidation, rather than the acquisition, phase of extinction learning due to our observation that sign-tracking behavior shows within-session extinction but impaired recall of extinction learning. Third, although doses were based on extinction studies in the existing literature, it is possible that different concentrations of the drugs used in the current study are necessary to facilitate the extinction of sign-tracking behavior. Further research is needed to determine the exact mechanisms underlying impaired extinction of sign-tracking behavior, investigate dose-response relationships of the drugs administered in the current study, and explore other pharmacological treatments targeting different signaling pathways (e.g., cannabinoid, opioid, noradrenergic, etc.) that might more effectively weaken resistance to extinction of sign-tracking behavior (Singewald et al., 2015).

## 5. Conclusion

Our results demonstrate that when reward-related cues are attributed with incentive-motivational value they become enduringly resistant to extinction and more prone to spontaneous recovery. Accordingly, current models of extinction should be updated to support individual variation in extinction of different CRs in response to the same CS. In addition, our results provide insight into several translationally relevant questions regarding the poor efficacy of cue-exposure therapy for addiction (Kantak and Nic Dhonnchadha, 2011). For example, the poor efficacy of cue-exposure therapy may not solely be the result of drug-induced neurocognitive impairments in drug-addicted patients, but rather a fundamental difference in the extinction sensitivity of cues imbued with incentive-motivational properties. Moreover, and based upon the results of the present study, caution should be taken when broadly applying research results from studies of Pavlovian fear conditioning to appetitive processes, particularly pharmacological investigations using cognitive enhancers. In the future, studies of appetitive Pavlovian extinction would benefit from the incorporation of PCA procedures to identify individual differences in the attribution of incentive-motivational value.

## Acknowledgements

We would like to thank Dr. Terry Robinson for material support of a portion of these studies and for contributions to the initial study design. This work was funded by the University of Michigan Department of Psychiatry (U032826 [JDM]) and Rackham Graduate School (Predoctoral Fellowship, CJF), the Department of Defense (DoD) National Defense Science and Engineering Graduate (NDSEG) Fellowship (CJF), the Brain & Behavior Research Foundation (NARSAD 20829 [JDM]), and the National Institute on Drug Abuse (NIDA; K08 DA037912-01 [JDM]; T32 DA007281 [CJF]).

